## Supplementary Information for "Random Matrix Theory-guided sparse PCA for single-cell RNA-seq data"

Victor Chardès*

Center for Computational Biology, Flatiron Institute, New York, NY, USA, 10010

#### A Almost sure limits of outlier eigenvalues in the separable covariance model

We can rewrite the system of equation for $g_1$ and $g_2$ in the form of a single equation, with $g_2(z)$ the unique solution $g_2$ of $F(z, g_2) = 0$ where $F$ reads

$$F(z, g_2) = g_2 - \int \frac{t\rho_A(t)}{-z\left(1 + tq\int \frac{t\rho_B(t)}{-z(1+tg_2)}dt\right)}dt. \tag{S1}$$

With the knowledge of $\rho_A$ and $\rho_B$, assuming we can exactly solve this equation, we have access to $m(z)$, and by the inversion formula to $\rho_S$ and its support.

##### A.1 Spiked separable covariance model

We consider here that the signal originates in the covariance only, such that $P = 0$. The question we address here is the following: given an eigenvalue $\alpha$ of $B$ such that $\alpha \notin \text{supp}\rho_B$, how is this eigenvalue reflected in the spectrum of $S$? To our knowledge, only partial results for this are available in the literature [1]. The eigenvalue $\alpha$ will give rise to outlier eigenvalues $\lambda \notin \text{supp}\rho_S$ satisfying the following equation

$$g_2(\lambda) = -1/\alpha. \tag{S2}$$

This equation for $\lambda$ is to be solved outside of $\text{supp}\rho_S$, and if there is no solution then it means that $\lambda$ falls on one of the edges $\rho_S$. We note that this result has only be proven for eigenvalues $\lambda$ above the rightmost edge of the spectrum [1], but numerical investigation (see Fig. S1) indicate that this result is more general. When there is a solution, there is no guarantee as to whether it is unique, and this equation can have multiple solutions, as also shown in Fig. S1. When $A$ is a low-rank deformation of the identity, i.e. $\rho_A(t) = \delta(t-1)$, Eq. S2 reduces to the classical result for spiked covariance matrices:

$$\underline{m}(\lambda) = -1/\alpha, \tag{S3}$$

for $\lambda \notin \text{supp}\rho_S$. In this case, because $\underline{m}(z)$ admits a functional inverse outside of $\text{supp}\rho_S$ [2], if there is a solution it is unique, such that one eigenvalue $\alpha$ gives rise to at most one outlier eigenvalue. In the case where $\alpha$ is a spike eigenvalue of $A$ rather than $B$, we can replace $g_2$ by $g_1$ in Eq. S2 [1].

##### A.2 Information-plus-noise model with separable covariance

We consider now the information-plus-noise model with separable covariance. The mean $P > 0$ is a low-rank matrix. To have a non-trivial limit $n \to \infty$ with $q = p/n$ fixed, the singular values of $P$ need to scale as $\sqrt{n}$. We denote such a singular value $\sqrt{\theta n}$ with $\theta = O(1)$. For this model, the question we address is the following: how is $\theta$ reflected in the spectrum of $S$? Relating $\theta$ to the eigenvalues of $S$ requires

*

the additionnal assumption that the matrices of left and right eigenvectors of $P$ are chosen uniformly at random in the space of orthogonal matrices [3]. In this case $\theta$ gives rise to eigenvalues solution of the equation

$$\lambda \underline{m}(\lambda) m(\lambda) = \frac{1}{\theta}, \tag{S4}$$

with $\lambda \notin \text{supp}\rho_S$ [3; 4]. The transform $z \mapsto zm(z)\underline{m}(z)$ is referred as the $D$-transform, and we have no guarantee that a functional inverse exists outside the support of $\rho_S$. For this reason, this equation may have multiple solutions. It was recently shown that in the case where $\rho_A(t) = \delta(t-1)$, the assumption about the distribution of the eigenvectors of $P$ can be dropped and any eigenvalue $\alpha$ of $A + P^T P/n$ outside the support of $\rho_A$ gives rise to outlier eigenvalues in the spectrum of $S$ that are solutions of equation Eq. S3. This result links the information-plus-noise model with the spiked covariance model: when $\rho_A(t) = \delta(t-1)$, irrespective of the model, any spike eigenvalue of $\mathbb{E}[S]$ will give rise to a single outlier eigenvalue satisfying criterion Eq. (S3).

We can rationalize this by verifying that, starting from Eq. S3, we can recover Eq. (S4) when the matrices of eigenvectors of $P$ are chosen at random in the space of orthogonal matrices. Since $\rho_A(t) = \delta(t-1)$, the functional inverse of $\underline{m}(z)$ is also explicit [2]. Using this inverse, along with $m(z) = (q^{-1}-1)/z + q^{-1}\underline{m}(z)$, we have

$$\lambda m(\lambda)\underline{m}(\lambda) = -\int \frac{\rho_B(t)}{t + 1/\underline{m}(\lambda)} dt = -m_B(-1/\underline{m}(\lambda)). \tag{S5}$$

We recognize $m_B$ the Stieljes transform of $\rho_B$. Because of the assumption on the eigenvectors, we can use a result on low-rank perturbations of symmetric random matrices stating that $\theta$ gives rise to outlier eigenvalues $\alpha$ in $\mathbb{E}[S]$ which are solutions of [5]:

$$m_B(\alpha) = -1/\theta. \tag{S6}$$

Combined with the previous equation and Eq. (S3), this allows us to recover Eq. (S4). In particular, while Eq. (S4) can have multiple solutions for $\lambda$, it's now Eq. (S6) that can have multiple solutions for $\alpha$. In this sense, the result relating outlier eigenvalues of $\mathbb{E}[S]$ to those of $S$ is more general (but not as insightful) than the information-plus-noise equation Eq. (S4) which requires an additional assumption on the eigenvectors of $P$.

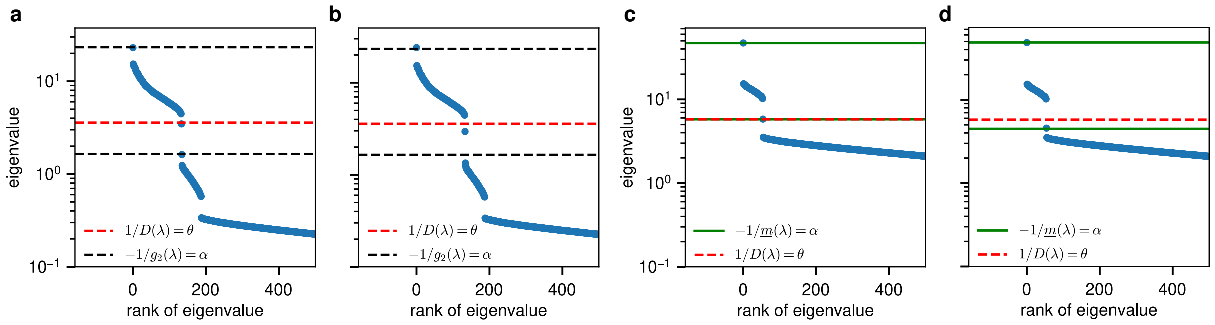


Figure S1: **Outlier eigenvalues for mixtures of information-plus-noise and separable spiked covariance models**. In this figure we use $n = 3180$ and $p = 3990$. We define a vector $u_1 \in \mathbb{R}^p$ with $u_{1,1} \approx 0.24$, $u_{1,2} \approx 0.97$, and all other entries zero. For the *correlated mixture*, we define a vector $u_2 \in \mathbb{R}^p$ with $u_{2,1} \approx 0.92$, $u_{2,2} \approx 0.39$, and all other entries zero, while for *independent* mixtures $u_2$ has entries chosen uniformly at random. We also define a vector $u_3 \in \mathbb{R}^n$ with entries chosen uniformly at random. All vectors are normalized. The matrices $A$ and $B$ are diagonal. We take $B$ with 60% of its entries equal to 12 and the rest equal to 1. The low-rank signals are defined as $Q = 45 u_1 u_1^T$ and $P = 2.5 u_3 u_2^T$. The data is generated as $X = A^{1/2} Y (B+Q)^{1/2} + P$ with entries of $Y$ being independent standard normal variables. The same realization $Y$ is used across all four subfigures. **a, b,** 30% of the entries of $A$ are set to 8 and the rest to 0.1. The signal eigenvalues $\alpha$ are computed as the isolated eigenvalues of $(B+Q)$. **c, d,** $A$ is the identity matrix. The signal eigenvalues $\alpha$ are computed as the isolated eigenvalues of $\mathbb{E}[S] = I + Q + P^T P/n$. **a, c.** Independent mixture. **b, d.** Correlated mixture.

#### A.3 Determination of the support

The previous equations are only usable if we have exact knowledge of the support of $\rho_S$. However, besides the case $\rho_A(t) = \rho_B(t) = \delta(t-1)$, analytical expressions for the Stieljes transform $m(z)$ and its associated density function $\rho_S$ are not known, and so isn't its support. It was shown that the edges of the support of $\rho_S$, $e_1 > ... > e_K \in \mathbb{R}^+$ of $\rho_S$ can be determined as the real solutions $(x, g_2) = (e_k, g_2(e_k))$ of this system of equation [1; 6]:

$$F(x, g_2) = 0 \text{ and } \frac{\partial F}{\partial g_2}(x, g_2) = 0. \tag{S7}$$

One can derive a more handy criterion by relating the support to the sign of the derivative $\partial F/\partial g_2$. In particular, it can be shown that any real solutions $(x, g_2)$ to the equation $F(x, g_2) = 0$ with $\partial F/\partial g_2(x, g_2) > 0$ verifies $x \notin \text{supp}\rho_S$ [1; 6]. This allows us to disregard the condition $\lambda \notin \text{supp}\rho_S$ in all the previous equations, solve them for $\lambda \in \mathbb{R}^+$, and check the sign of the derivative $\partial F/\partial g_2(\lambda, g_2(\lambda)) > 0$. In particular, given a solution $(\lambda, g_2(\lambda))$ to the outlier eigenvalue equation (for the spiked covariance model or the information-plus-noise model), this criterion can be rewritten as:

$$1 - \frac{q}{(\lambda g_2(\lambda) g_1(\lambda))^2} \int \frac{t^2 \rho_B(t)}{(t + 1/g_2(\lambda))^2} dt \int \frac{t^2 \rho_A(t)}{(t + 1/g_1(\lambda))^2} dt > 0. \tag{S8}$$

In the case $\rho_A = \delta(t-1)$, this criterion simplifies, and we recover the one for the spiked covariance model [2], where $\psi$ denotes the functional inverse of $-1/\underline{m}(\lambda)$:

$$\psi'(\alpha) > 0 \text{ with } \psi(\alpha) = \alpha + q\alpha \int \frac{t\rho_B(t)}{\alpha - t} dt. \tag{S9}$$

Any outlier eigenvalue associated with a spike eigenvalue $\alpha$ that does not satisfy the criterion will have as almost sure limit a point at an edge of the support of $\rho_S$. For the sake of this paper, it means that it can't be distinguished from the bulk. We note however that by leveraging additional results from RMT this could *in theory* be done [7].

#### A.4 Choice of a model

The inverse problem at hand is the following: given outlier eigenvalues observed in $S$, we want to infer the associated signal eigenvalue in $\mathbb{E}[S]$. For this, we need to assume an underlying random matrix model, but we would like to do so with minimal assumptions. We have the following choices:

1. Independent mixture of spiked separable covariance and information-plus-noise models: right and left eigenvectors of $P$ are chosen uniformly at random in the space of orthogonal matrices and $\rho_A \neq \delta(t-1)$. In this case, both Eq. S4 and Eq. S3 hold, as shown in Fig. S1a.

2. Correlated mixture of spiked separable covariance and information-plus-noise models: $P$ and $B$ are not independent and $\rho_A \neq \delta(t-1)$. Eq. S4 and Eq. S3 can't be reliably used to predict the position of outlier eigenvalues, as shown in Fig. S1c.

3. Correlated or independent mixture of left-whitened spiked covariance and information-plus-noise models: $P$ and $B$ may or may not be independent and $\rho_A = \delta(t-1)$. In this case, Eq. S3 relates all outlier eigenvalues of $S$ to those of $\mathbb{E}[S]$, as shown in Fig. S1c,d.

In the first model, we don't have a single mapping relating eigenvalues of $S$ to those of $\mathbb{E}[S]$. Without additional information like fluctuations around the almost sure limits of outliers, we cannot choose between the information-plus-noise or the spiked covariance models. In the second model, to our knowledge we don't have a consistent mapping relating all outlier eigenvalues to signal eigenvalues. Finally, in the third model we do not need any assumption on the left and right eigenvectors of $P$, and we have a unique formula to relate the outlier eigenvalues of $S$ to those of $\mathbb{E}[S]$. To use this model it is necessary to first whiten the cell-cell covariance, i.e. it is only applicable to $X \leftarrow A^{-1/2}X$.

### B Reproducibility

For the sake of reproducibility, we provide flow charts describing the quality control and feature selection steps, as well as the design of the benchmark. These flow charts are shown in Fig. S2, Fig. S3-Fig. S5.

##### B.1 Quality control and feature selection, Fig. S2

The default gene selection method is `flavor='seurat'` from the `scanpy` package [8]. When using `flavor='seurat_v3'`, the library-size and log-normalization steps are removed before selecting the set of genes, as recommended by the method. Each pipeline shown in Fig. S2 returns count data and serves as the basis for all experiments in this paper.

##### B.2 Benchmark design, Fig. S3-S5

Each horizontal branch from the main (left-side) pipeline corresponds to an independent copy of the data. Each copy is then processed through the different downstream pipelines. This design ensures that all methods receive exactly the same data, with identical features and cells. The red arrows indicate that the set of genes selected at earlier steps (quality control and highly variable gene filters) is reused across pipelines. This ensures that the full dataset is processed using the same set of genes as the subsampled dataset. Otherwise, the set of genes selected on the full dataset would differ substantially from that obtained on the subsample. These precautions are necessary to guarantee that each method is evaluated fairly and not biased by differences in gene selection.

The final step of each pipeline is a low-dimensional embedding of the cells. For scVI and DCA, each dataset produces two distinct low-dimensional embeddings (PCA and latent) from the same data copy, though the latent embedding is not shown in the flow chart. The parameters displayed correspond to one realization of the trials and the number of components used in the PCA steps may vary slightly. Since DCA and scVI are hyperoptimized, the parameter set for these methods may differ from one trial to another.

**Algorithm S1** FISTA sparse PCA

**input:** sample covariance matrix $S$, leading eigenvectors $V$ (ordered columnwise)
**output:** sparse loading matrix $W$
Operations max, abs, $\times$ and sign are performed entry-wise
$p \leftarrow 1/20,\ q \leftarrow 1,\ r \leftarrow 4$
$\gamma \leftarrow 1/(2\lambda_{\max}(S))$, where $\lambda_{\max}(S)$ denotes the max eigenvalue of $S$
$W_0 \leftarrow V,\ Y_0 \leftarrow V,\ t_0 \leftarrow 1$
**while** stopping criterion not reached **do**
  $Z \leftarrow Y_k + 2\gamma S Y_k$
  $Z \leftarrow \max\big(\text{abs}(Z) - \lambda\gamma,\, 0\big) \times \text{sign}(Z)$
  $Z \leftarrow \textsc{orthogonalize}(Z)$ ▷ e.g. Löwdin or Gram-Schmidt
  $t_{k+1} \leftarrow (p + \sqrt{q + r t_k^2})/2$
  $Y_{k+1} \leftarrow Z + \dfrac{t_k - 1}{t_{k+1}}(Z - W_k)$
  $W_{k+1} \leftarrow Z$
  $k \leftarrow k + 1$
**end while**
**return** $W_k$

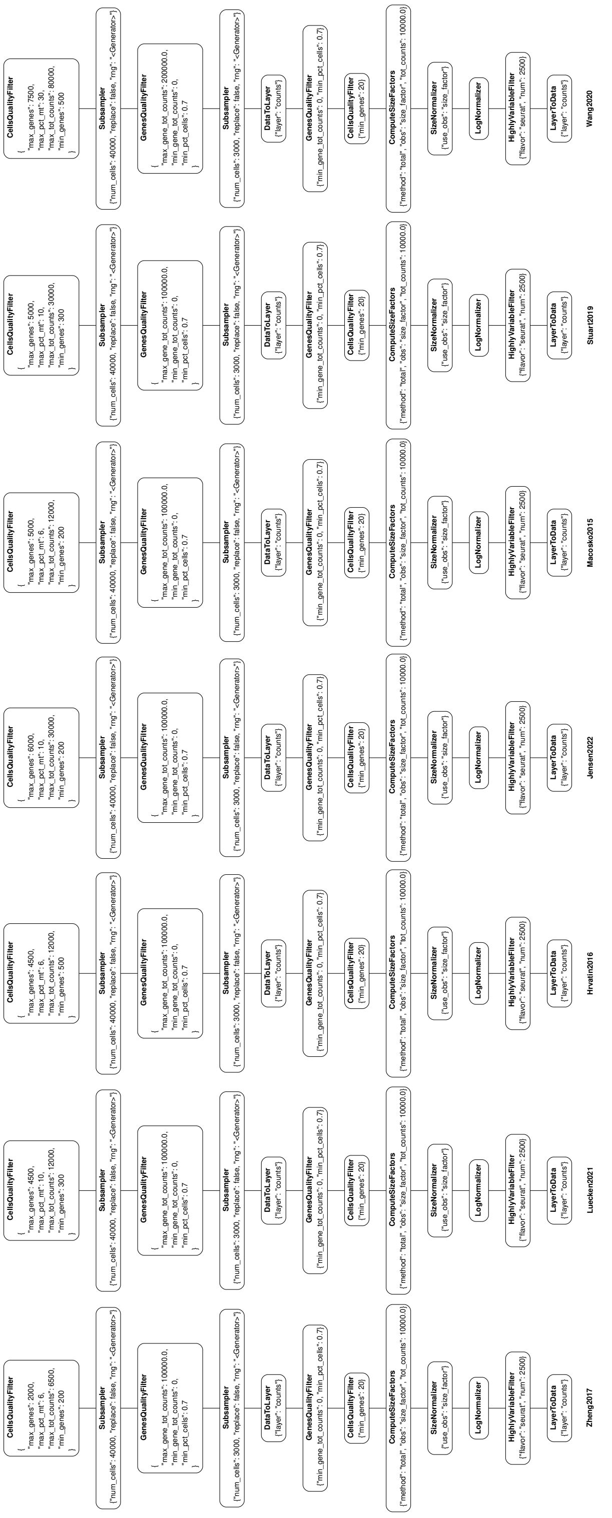

Figure S2: **Quality control and feature selection pipelines**. This figure is rotated 90 degrees. Each pipeline is read from top to bottom. All quality filters are applied as strict inequalities on the specified thresholds.

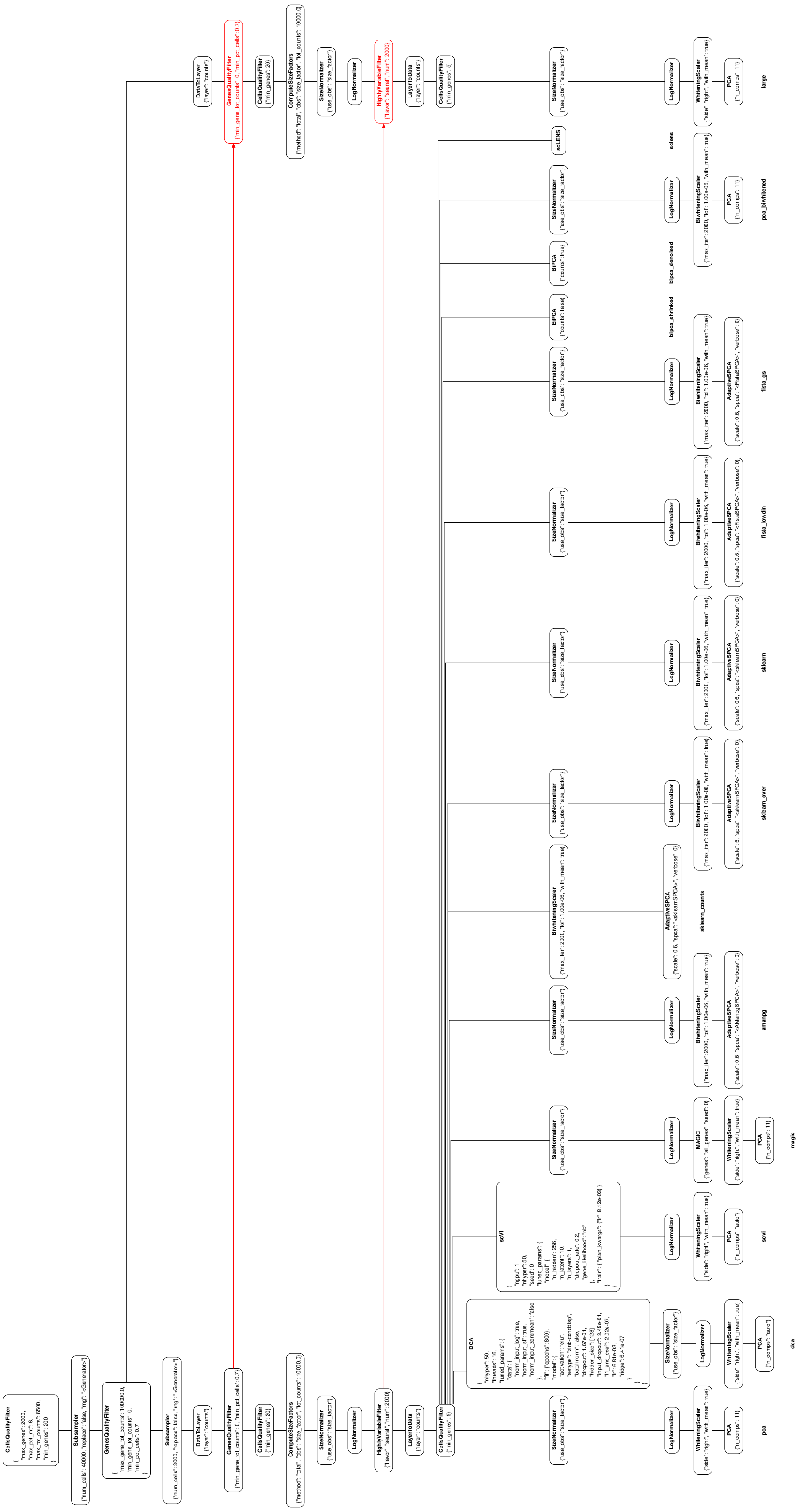

Figure S3: **Design of the benchmark for Zheng2017**. This figure is rotated 90 degrees. The flow chart is read from top to bottom.

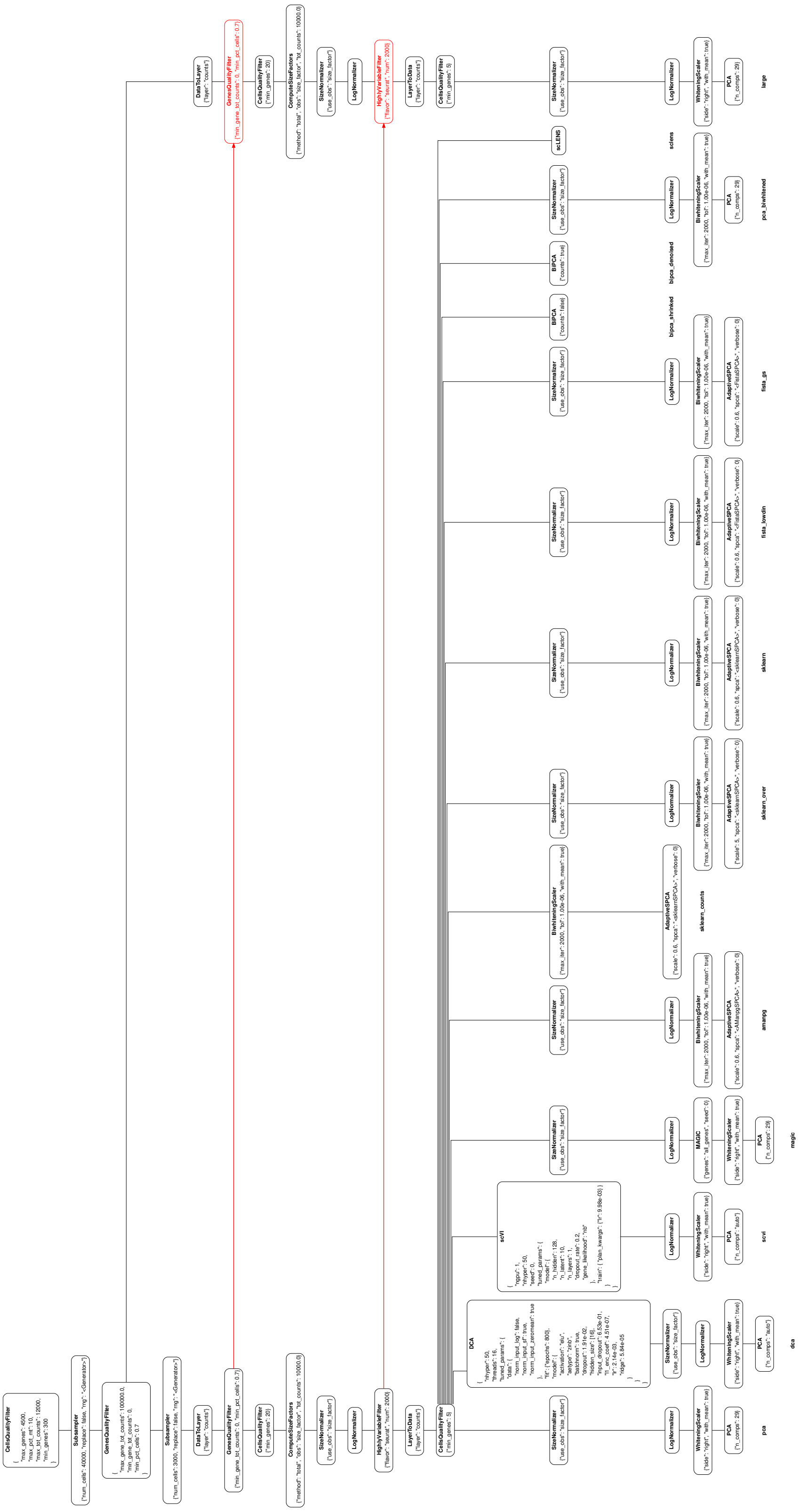

Figure S4: **Design of the benchmark for Luecken2021**. This figure is rotated 90 degrees. The flow chart is read from top to bottom.

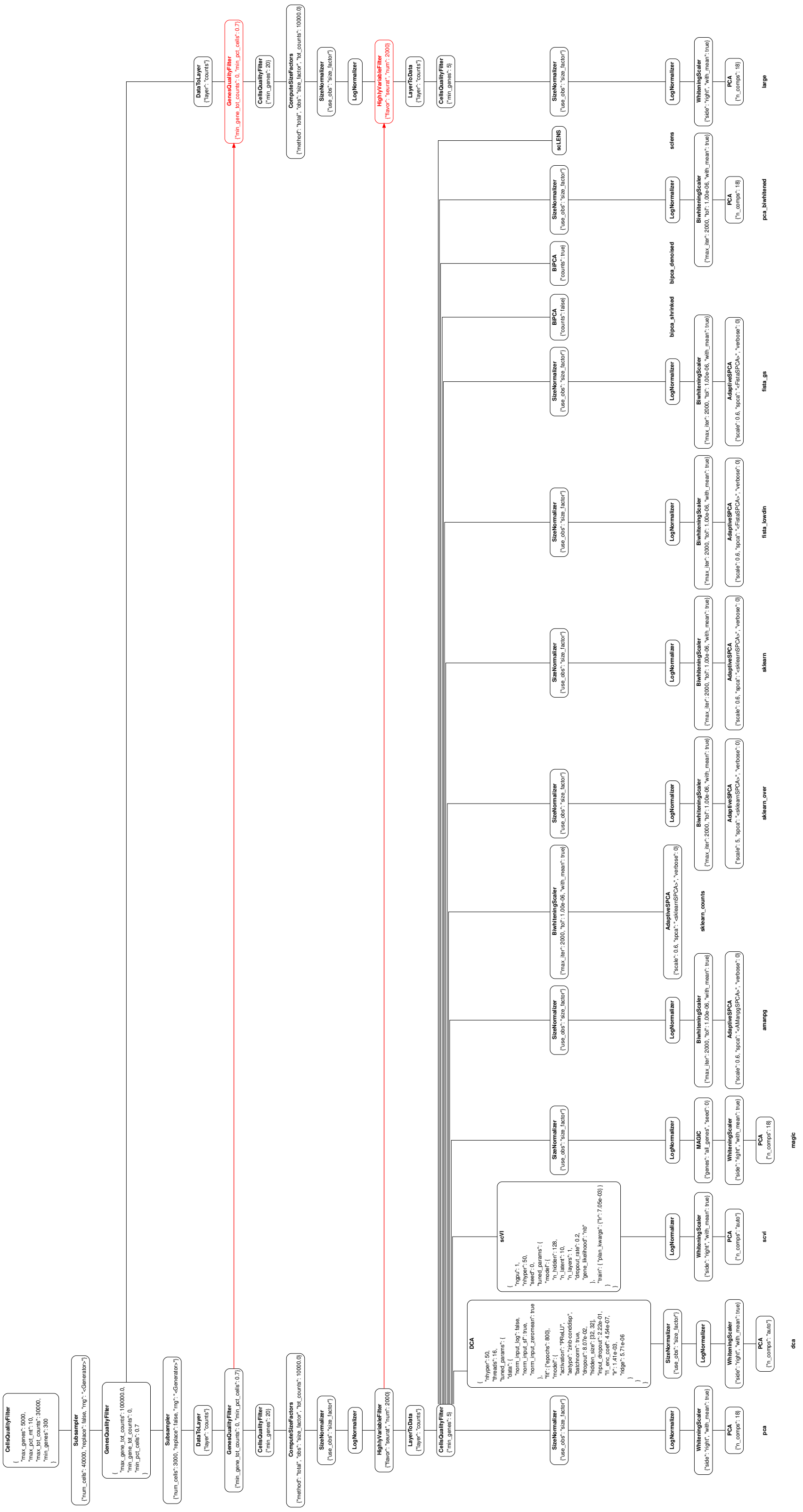

Figure S5: **Design of the benchmark for Stuart2019**. This figure is rotated 90 degrees. The flow chart is read from top to bottom.

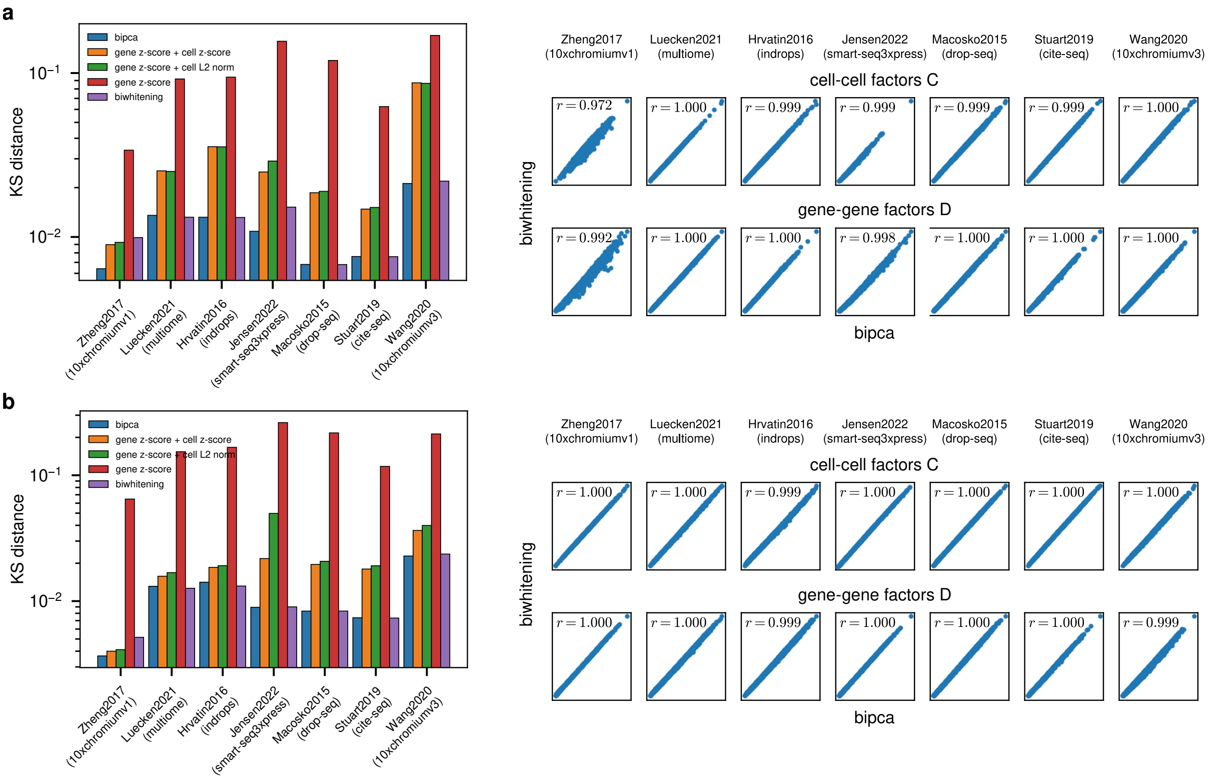


Figure S6: **Comparison of Biwhitening and BiPCA**. We compare the bi-proportional scaling from the BiPCA package with our biwhitening approach on count data [9] Both methods perform on par, yielding almost identical biwhitening factors. **a.** Results for 2500 highly variable genes; **b.** results for 10000 highly variable genes.

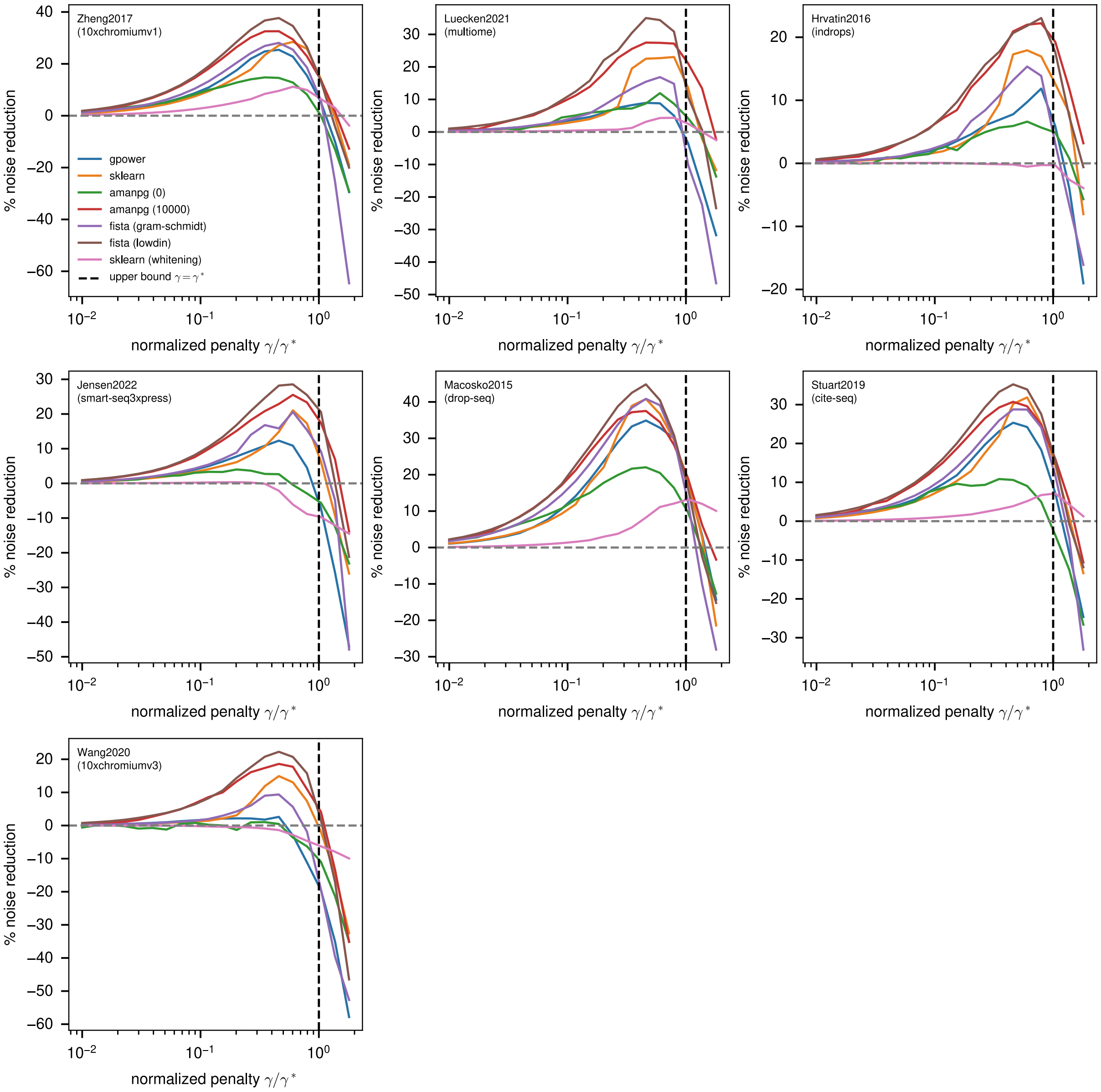


Figure S7: **Principal subspace reconstruction**. Noise reduction after RMT-guided sparse PCA for all datasets and all sparse PCA algorithm. The results discussed in the main text generalize to all datasets.

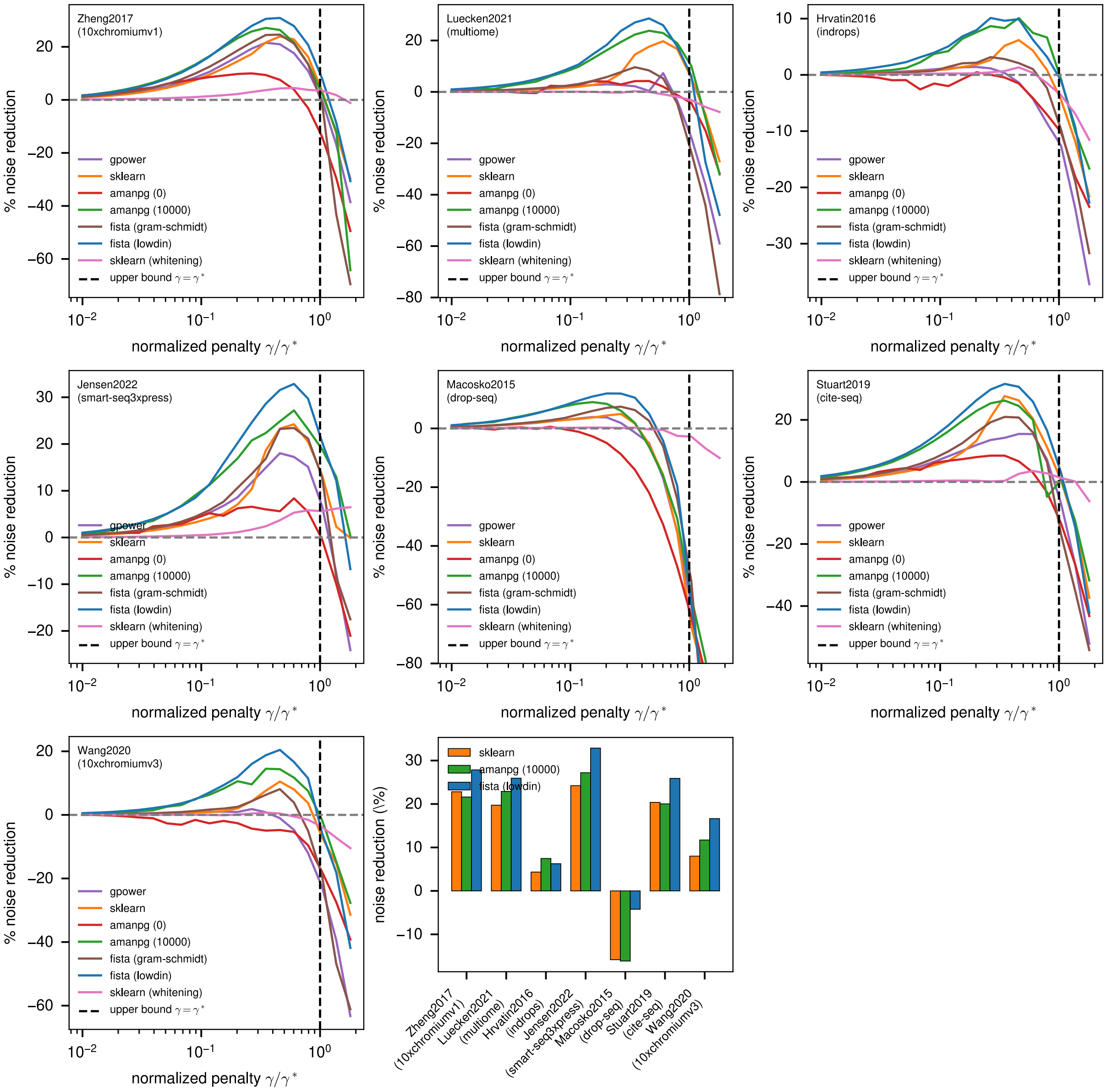


Figure S8: **Principal subspace reconstruction with different gene sets**. Noise reduction as a function of the penalty parameter for all datasets and all sparse PCA methods using 2000 highly variable genes selected with the parameter `flavor='seurat_v3'` in the `scanpy` package [8], see Fig. S2.

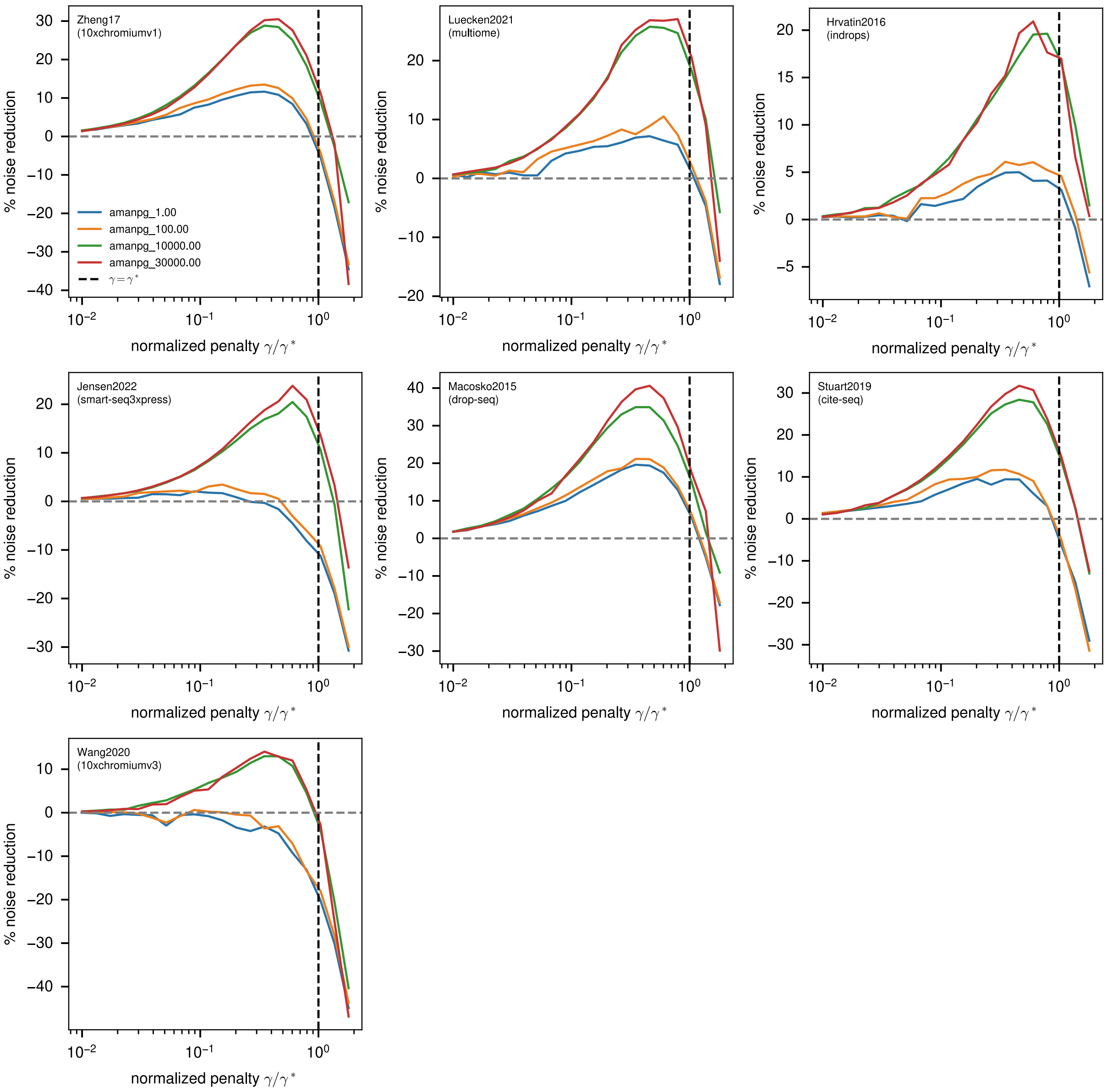


Figure S9: **AManPG algorithm with different $L_2$ penalties**. Noise reduction as a function of the penalty parameter for all datasets using AManPG with $L_2$ penalties $\eta = 1, 10^2, 10^4, 3 \cdot 10^4$. A very large penalty yields the best performance.

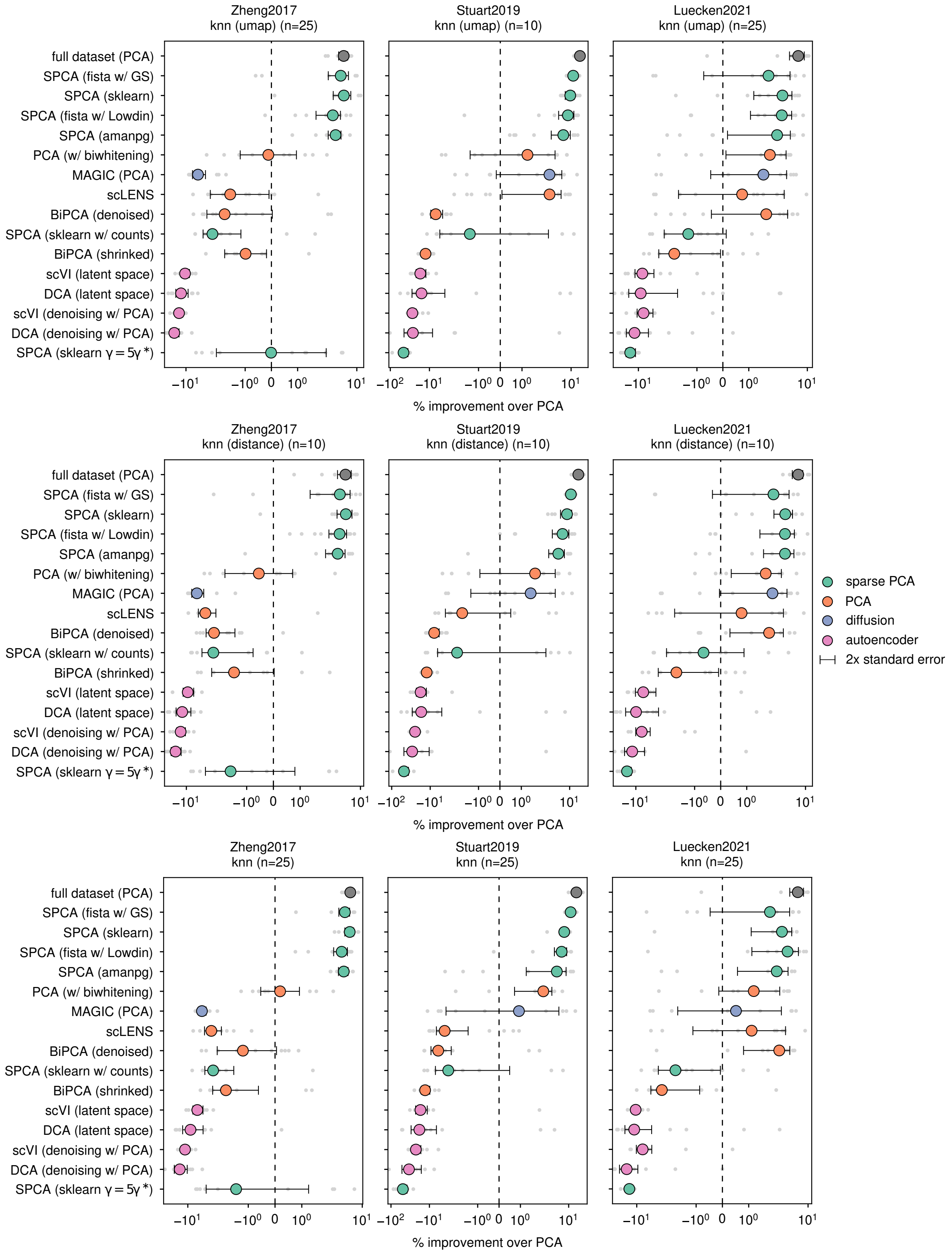


Figure S10: $k$**-NN classification performance**. Cell type classification performance as evaluated with bagged predictors of 30 classifiers, measured as a an improvement with respect to the results obtained with PCA. Top row: $n = 25$ neighbors with umap weights. Middle row: $n = 10$ neighbors with inverse distance weights. Bottom row: $n = 25$ neighbors with constant weights.

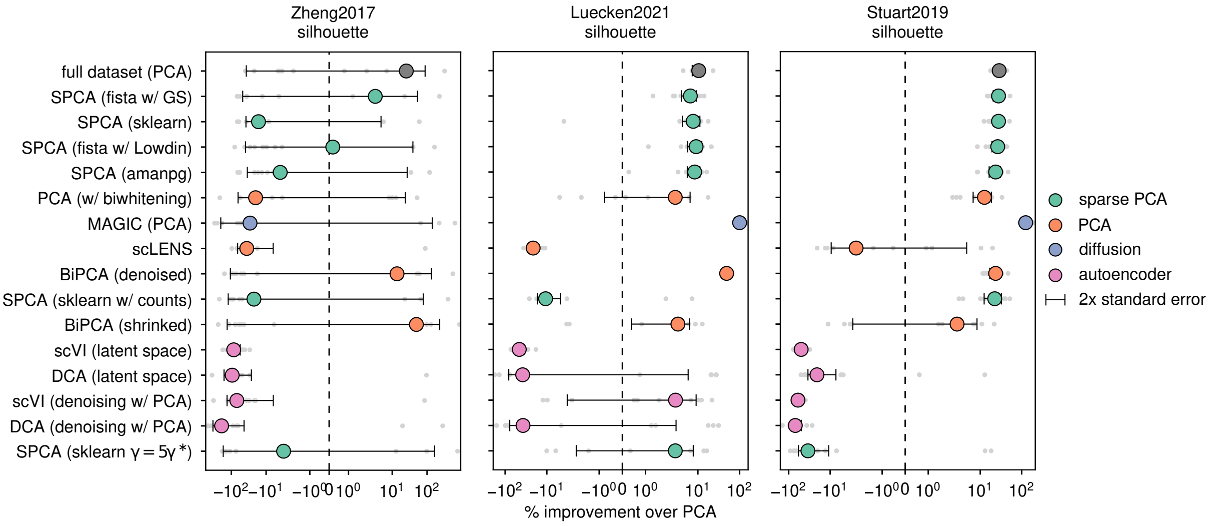


Figure S11: **Average silhouette score**. Average silhouette score for the ground-truth cell-type annotation in the projected lower-dimensional spaces. The Zheng2017 dataset is particularly challenging, with highly mixed annotations. For this dataset, the average silhouette score of PCA is close to zero, leading to non-significant findings. Both MAGIC and BiPCA perform very well on the two other datasets, despite performing poorly on the $k$-NN benchmark.

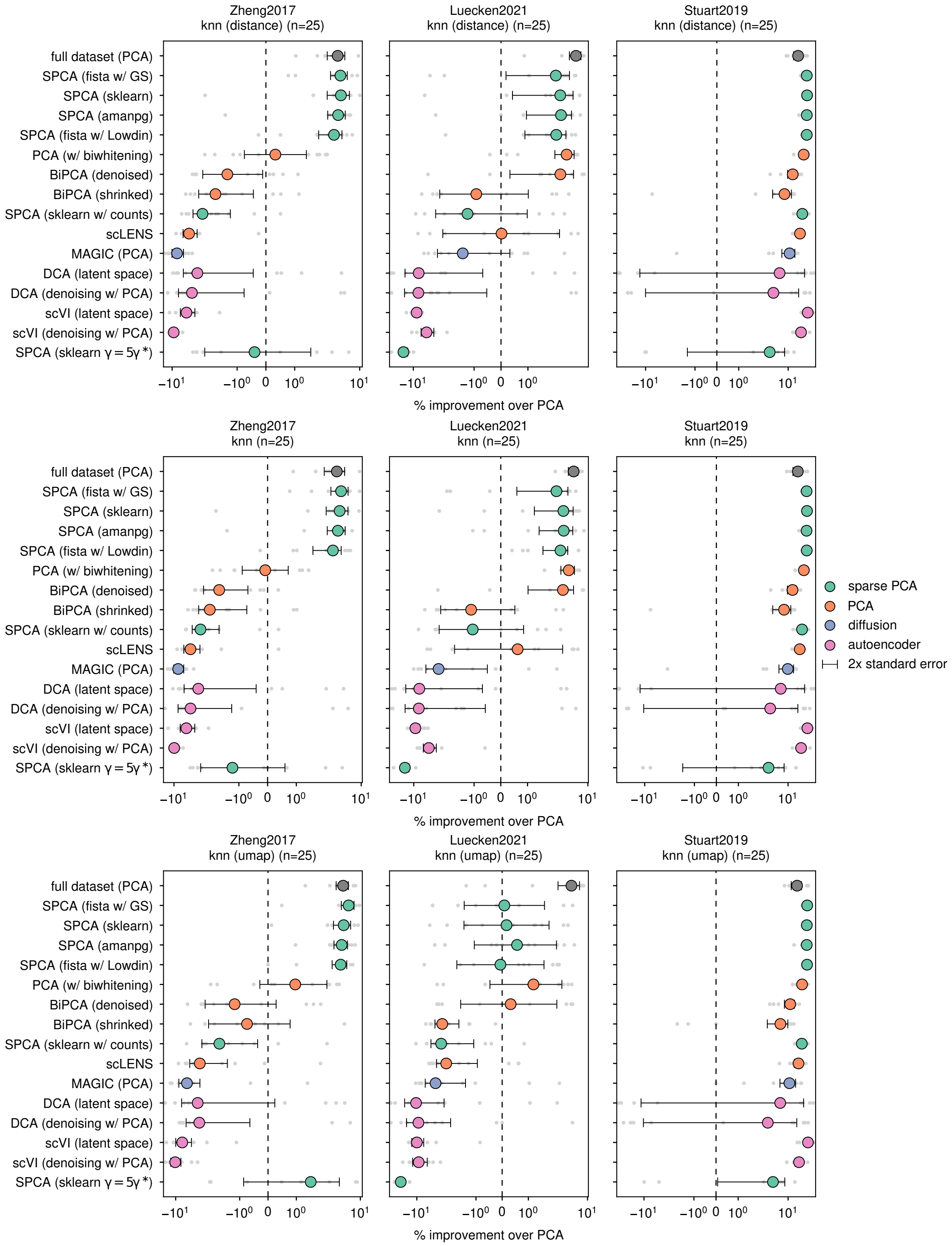


Figure S12: $k$**-NN classification performance with different gene sets**. Cell type classification performance evaluated using the parameter `flavor='seurat_v3'` in the `scanpy` package [8], see Fig. S2.

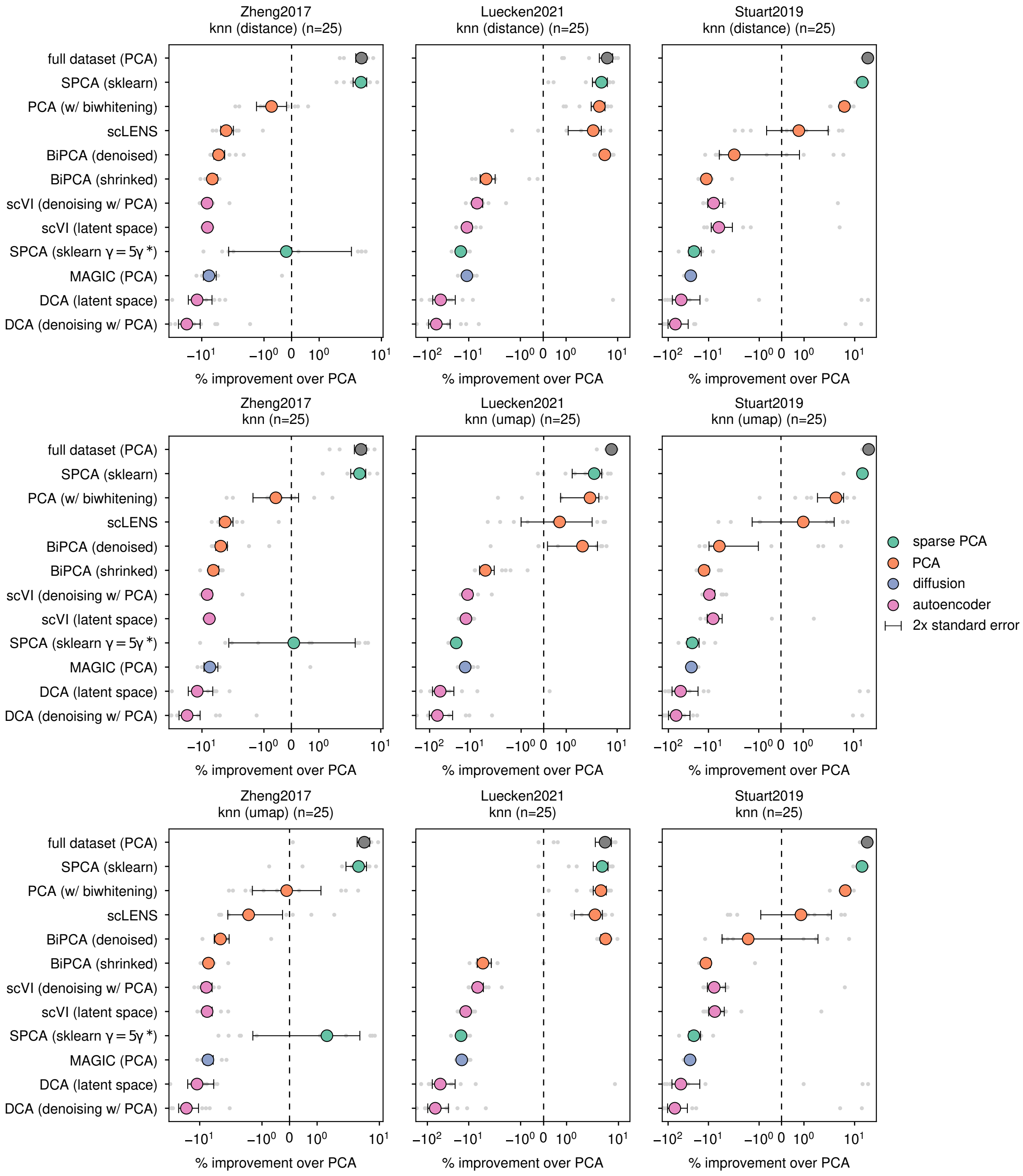


Figure S13: **$k$-NN classification performance with** 10000 **highly variable genes**. Cell type classification performance evaluated Fig. S10, but using $n = 10000$ highly variable genes.
